# Dose-dependent growth and gene expression in marine prokaryotes in response to glacial and non-glacial iron-rich colloids from Kerguelen Island (Southern Ocean)

**DOI:** 10.64898/2026.10.28.761898

**Authors:** Carla Pereira Garcia, Urania Christaki, Rhea Thoppil, Audrey Guénueguès, Philippe Catala, Olivier Crispi, Stéphane Blain, Ingrid Obernosterer

**Author notes:** Carla Pereira Garcia Urania Christaki Rhea Thoppil, Audrey Guénueguès Philippe Catala, Olivier Crispi Stephane Blain.

## Abstract

Iron (Fe) plays a central role in regulating prokaryotic carbon (C) metabolism through complex processes. In Fe-limited limited ocean regions, new sources can influence prokaryotic activity thereby modulating the microbially mediated cycling of carbon. However, while accelerated glacial melting delivers substantial quantities of Fe-rich material to the ocean, the bioavailability of glacial Fe remains poorly understood. In this study, we investigated the response of marine prokaryotes to Fe contained in the colloidal fraction (20–200 nm size fraction) originating from the Cook Ice Cap on Kerguelen Island (Southern Ocean). We assessed growth and gene transcriptional responses to dose-dependent additions of Fe-rich colloids from two sources, glacial meltwater and a non-glacial origin. The two colloid types resulted in dose-dependent, but contrasting growth responses. Increasing additions of glacial colloids resulted in a reduction in prokaryotic growth and the opposite pattern was observed with non-glacial colloids. Transcriptional profiles revealed an increasing trend in Fe-limitation, as evidenced by transcripts of the glyoxylate shunt, as a function of the addition of either colloid type. We further observed a dose-dependent increase in transcripts of aromatic compound degradation only in the non-glacial colloid incubations, suggesting that these Fe-rich colloids could also provide C. These results point to the key role of C regulating prokaryotic growth in response to Fe-rich colloids. The quantitative transcriptional responses observed provide insights to the nutritional status of prokaryotes and to compounds and pathways involved in processing Fe-rich colloids.

## Introduction

Organic carbon (C) and iron (Fe) are essential for prokaryotic metabolism, but both can be co-limiting heterotrophic activity and growth in different ocean regions (Church et al. 2000; Kirchman et al. 2003; Obernosterer et al. 2015; Baltar et al. 2018; Manck et al. 2024). Phytoplankton can play a dual role in this context. They supply heterotrophs with labile organic substrates, but phototrophs can also compete for Fe (III) inorganic ions (Fe’) a highly labile form of Fe (Fourquez et al. 2015). Thus, Fe bioavailability regulates phytoplankton-derived organic matter production, which in turn affects the degree of Fe- and C (co-) limitation of prokaryotes. To access non-directly available forms of Fe, such as those bound to organic ligands (Gledhill and Buck 2012) prokaryotes can produce siderophores, low molecular weight compounds, that can strongly chelate Fe, rendering complex Fe sources bioavailable (Sandy and Butler 2009) . However, siderophore production is energy-intensive adding another layer of complexity to prokaryote-phytoplankton interactions.

Novel Fe sources can alleviate limitation at varying spatial and temporal scales, but the chemical forms of Fe introduced to the ocean and the potential role of prokaryotes in accessing them are poorly understood. For example, atmospheric deposition or hydrothermal vents contain Fe embedded in mineral-type particles (Schroth et al. 2009; Bühring et al. 2025), requiring transformations prior its utilization. The accelerated melting of ice sheets and glaciers present a quantitatively important source of Fe for polar oceans (Van Der Merwe et al. 2019; Hawkings et al. 2026). Glacial erosion releases Fe into the ocean primarily as low-solubility mineral particles (Schlosser et al. 2012). While a large fraction of these Fe-rich particles rapidly aggregates and sinks upon entering seawater, a portion remains in the nanometer size-fraction (Hawkings et al. 2014). This colloidal Fe, which often exists as Fe oxyhydroxide or mineral-associated forms, can be transported from coastal to the open ocean regions. However, its bioavailability remains unclear. A recent study reports that glacial colloids stimulated the growth of specific marine prokaryotic taxa and increased the abundance siderophore production and transport genes as compared to colloids of non-glacial origin (Thoppil 2025). This response based on the genetic potential, raises the question of whether transcriptional profiles provide more detailed insights to the mechanisms and pathways involved in the processing of colloidal Fe.

Metatranscriptomics is now widely used in environmental studies, yet connecting gene expression profiles to microbial metabolism in a quantitative manner remains a major challenge (Saito et al. 2024). Gene expression is regulated by multiple environmental cues and both expression patterns and levels can be taxon-specific (Gifford et al. 2013; Peoples et al. 2026) rendering it difficult to obtain quantitative estimates from transcriptional activities within complex microbial communities. To address this, transcriptional markers identified for bacterial model organisms under controlled experimental conditions were proposed to interpret *in situ* gene expression (Gifford et al. 2016). Using ship-board experiments with gradual Fe-availability modifications revealed the co-existence of *Synechococcus* ecotypes adapted to different Fe- regimes and the expression of specific genes provided insights on Fe-limitation of these taxa *in situ* (Gilbert et al. 2022). These studies provide insights on how transcriptional activity can inform on the nutritional status of taxa within microbial communities.

The goal of this study was to determine how marine prokaryotes respond to Fe in the form of colloids of glacial or non-glacial origin. To achieve this, we investigated the bulk and metatranscriptomic responses to increasing additions of Fe-rich colloids in incubation experiments. In addition to this quantitative approach, we analyzed gene expression using three metrics that are transcripts per million (TPM), transcripts per liter (TPL, Satinsky et al. 2013) and transcripts per active cell (Salazar et al. 2019). Our findings demonstrate that dose additions of limiting nutrients in incubation experiments can induce quantitative transcriptional responses across all metrics. These responses provide insights to the nutritional status of prokaryotes and reveal the mechanisms underlying Fe and C co-limitation.

## Material and Methods

### Environmental context

Water samples for the collection of colloids were taken in two lakes on Kerguelen Island (Indian Sector of the Southern Ocean, 49°S, 70°E) during the project MARGO (December 2023-February 2024) (Fig. 1). The two lakes are geographically close (0.6 km) and have both a mineral-type catchment area, dominated by basalt, but differ with respect to the influence by the Cook Ice Cap (CIC) that is experiencing a dramatic wastage since 1960 (Verfaillie et al. 2021).

**Fig. 1.**
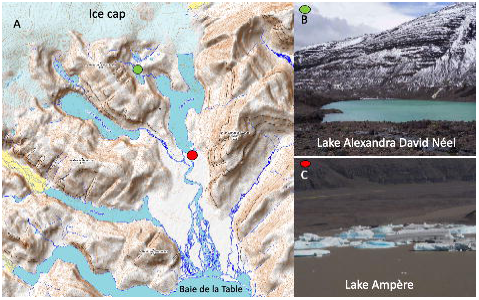
Map of the southern part of Kerguelen Island with the sites where the water for colloid collection was sampled are shown by colored dots (A). Pictures of Lake Alexandra David Néel (B) and Lake Ampère (C). From Thoppil et al. (2025).

Lake Ampère is directly connected with one of the glaciers (Glacier Ampère) of the CIC and therefore receives icebergs and meltwater influenced by glacial erosion. The outflow of Lake Ampère enters the coastal site *Baie de la Table*. By contrast, Lake Alexandra David Néel is located at 200 m altitude with no connection to the CIC and this lake has therefore no contemporary input of glacial meltwater.

For clarity, we will refer in the following to Lake Ampère as glacial lake and to Lake Alexandra David Néel as non-glacial lake. Water samples were collected at the surface using a 6L Niskin bottle that was deployed horizontally about 5 m from the coast. The Niskin bottle was closed from distance and pulled back to the shore, turned vertically and samples were then drawn from the spigot. For the environmental characterization of these two freshwater systems, samples were taken for the inorganic nutrients nitrate (NO_3_ ^-^), phosphate (PO_4_ ^3-^), silicic acid Si(OH) and dissolved iron (DFe), and also for microbial cell abundances and community composition (16S rRNA gene sequencing) and the respective analyses are described in the Supporting Information.

### Preparation of colloids by ultrafiltration

Colloids were obtained from both lakes by ultrafiltration as described in Thoppil et al. (2025). Briefly, lake water was sequentially filtered through 0.8 µm and 0.2 µm pore size Polycarbonate (PC) filters and the < 0.2 µm filtrate was then passed through an ultrafiltration system (Pellicon, Millipore) equipped with 100 kilo Dalton (kDa) cassette filters. The Retentate of the UltraFiltration (RUF) contains colloids > 100 kDa, corresponding to approximately 20 nm. Our aim was to prepare colloidal concentrates of each lake that allowed the addition of equal volumes to obtain a given concentration of Fe in the respective incubations. We therefore adjusted the concentrations factors, based on the Fe concentrations determined in the respective colloidal fractions during a previous sampling campaign (Thoppil et al. 2025). The concentration factors were 22.2 and 88.8 for water from the glacial non-glacial lakes, respectively.

### Microbial incubation experiments

To determine the response of marine microbial communities to dose additions of colloids originating from the two different lakes, we performed short-term incubations (4 days) with a prokaryotic community from a High Nutrient Low Chlorophyll site located south of Kerguelen Island (52.54°S, 67.33°E). Seawater was collected at 10 m depth using a trace-metal clean GEO-FISH (Baudet et al. 2024). Concentrations of DFe and dissolved organic carbon were 0.3 nM and 62 µM, respectively, and concentrations of major nutrients were 24.1 µM for NO_3_ ^-^, 1.56 µM for PO_4_ ^3-^, and 9.3 µM for Si(OH) at the depth of sample collection. For the incubations, about 15L were filtered through 1 µm (Whatman Polycap PP filter cartidges) in a trace-metal clean lab and the < 1 µm filtered seawater was dispensed to 1L acid-cleaned Polycarbonate (PC) bottles (Supporting information Fig. S1). Increasing volumes of the either non-glacial or glacial retentates (5.56 mL, 17.78 mL and 55.56 mL) were added to 1L of < 1 µm filtered seawater. In another set of incubation bottles, we added Fe-EDTA with final concentrations of 10 nM, 32 nM and 100 nM. Unamended incubations (3 x 1L bottles) served as control. All PC bottles were placed in an indoor incubator and maintained in the dark at 4°C for 4 days. After this incubation period, the total volume was split into two subsamples of 500 mL. Each of the 500 mL was filtered onto PC filters (Millipore; 0.2 µm pore-size, 47 mm diameter), placed in a 1.8 mL cryotube, shock-frozen and stored at -80°C until extraction of RNA or DNA. The incubations were also subsampled for the enumeration of microbial abundances (prokaryotes and heterotrophic nanoflagellates) using previously described protocols (see Supporting Information).

### RNA and DNA extraction

RNA was extracted for all incubations, except for 1 unamended control, which was used for DNA extraction. In addition, DNA extracted from one sample collected at the HNLC site was used for metagenomic sequencing. DNA extraction was carried out as described previously and is detailed in the Supporting Information. RNA was extracted using the RNeasy PowerWater Kit (Qiagen, Hilden, Germany), following the manufacturer’s protocol with minor modifications to optimize the lysis of the cells and the preservation of the RNA. Prior to starting the protocol, the filter was frozen using liquid nitrogen and crushed with a mortar, the crushed pieces were added to the PowerWater Bead Pro Tube. As a first step in the manufacture’s protocol, we added the PW1 and ß-mercaptoethanol, additionally we added 66 µl of lysozyme at 20mg/ml and incubate it at 37°C for 45 minutes, followed by the addition of 14 µl of proteinase K at 20mg/ml and incubation at 55°C for 60 minutes. Final RNA concentrations were measured in the home laboratory using the QuantiFluor® RNA System. To enable quantification of absolute transcript abundances, internal standards were added at the initial step of the extraction. This approach ensured consistency and reliability in subsequent internal standard-based abundance calculations. Details on the selection of internal standards and the quantification approach are provided in the Supporting Information.

Metatranscriptomic and metagenomic samples were sequenced using an Illumina NovaSeq 6000 system with 2 × 150 bp chemistry at Fasteris SA, Inc. (Switzerland). Each adapter-trimmed sample yielded 30-82 million pairs of metatranscriptomic reads and 110-130 million pairs of metagenomic reads. The datasets can be accessed in the ENA repository under the PRJEB113620 and basic information about the reads can be found in Supporting Information Table S1.

### Bioinformatic pipeline

The raw metatranscriptomic sequences were used directly as input for the nf-core pipeline *metatdenovo* (A. Ewels et al. 2020; Lundin et al. 2025) in which the following versions of the databases were used: SILVA SSU and LSU 138.2 (Quast et al. 2012), GTDB r220 (Parks et al. 2022), NCBI nr database v.2024-02-07 (Goldfarb et al. 2025), KEGG database downloaded on 22-05-2025 (Kanehisa and Goto 2000). Briefly, the pipeline performed quality assessment and filtering of the RNA reads, an additional filtering step to remove remaining rRNA, uses megahit to create a *de novo* assembly with all the metatranscriptomic samples and then *prodigal* and/or *prokka* for gene (or transcript) prediction. Functional and taxonomical annotations for the predicted transcripts, as well as mapping and read counting to obtain their relative abundances are also included in the pipeline. Additionally, to further characterize the assemblies, we used the FeGenie tool (Garber et al. 2020) for Fe related genes (options --orfs --all_results) and dbCAN3 v.4.1.4 (Zheng et al. 2023), to annotate CAZymes and organic substrate transporters.

The raw metagenomic reads were filtered with fastp (Chen et al. 2018) and then assembled with megahit (Li et al. 2016)(options - – min-contig-len 1000). Prodigal was use to predict the open reading frames and prokka (Seemann 2014), kofamscan (Aramaki et al. 2019) and FeGenie (Garber et al. 2020) were used for further functional annotation. Due to technical complications (contigs were too short) with the use of FeGenie, the results from the metatranscriptomic de novo contigs needed to be validated. A blastn (Altschul 1997) was done between the positive results from FeGenie in the 2 metagenomic samples (considered true positive results) and the positive results from the metatranscriptomic samples and only hits with >95% identity and 100% query coverage were kept.

### Normalization approaches

We used three ways of normalizing metatranscriptomic sequences and the details on the calculations are provided in the Supporting Information. Transcripts per million (TPM) is a metric commonly used to assess the relative abundance of transcripts, taking into account the gene length and the sequencing depth of the sample (Wagner et al. 2012). Transcripts per liter (TPL) are obtained through the use of an internal standard added during the first step of the extraction to calculate the total transcripts of interest in the sample (Satinsky et al. 2013). A recent approach uses ten single copy genes to normalize the number of transcripts to the transcriptionally active cells (Sunagawa et al. 2013; Milanese et al. 2019; Salazar et al. 2019), and we refer here to this metric as transcript per cell (TPC).

### Fe-related gene database

To investigate possible relationships between the addition of colloids in a dose-dependent manner and the transcription of genes related to Fe, we compiled a list of genes in which Fe is needed either to assure the functionality of the protein or is part of the substrate of said protein. We created this list using proteins from FeGenie (Garber et al. 2020), all the metalloenzymes in which Fe is required (Metal-MACiE; (Andreini et al. 2009) as well as other known Fe-metalloproteins (Debeljak et al. 2019; Manck et al. 2024) (Supporting Information Table S2). To assess the transcriptional responses in the different treatments, we calculated whether individual transcripts presented an increasing trend. We defined a trend as at least a 5% increase between each addition of Fe in colloidal form or as Fe-EDTA for all metrics considered here. To assess whether transcripts related to Fe (Table S2) had a statistically higher probability for a trend as compared to transcripts not related to Fe, we performed a Chi-Square test, a frequency analysis, using the R package *stats* (RStudio Team 2020; R Core Team 2023). All the figures were computed using the R package *ggplot2* (Wickham 2016; RStudio Team 2020; R Core Team 2023).

## Results

### Environmental context in the non-glacial and glacial lakes

Among the abiotic parameters determined, concentrations of DFe revealed the most pronounced differences with 8.5-fold higher concentrations in the glacial (964 nM) as compared to the non-glacial lake (113 nM)(Table 1). DOC concentrations were overall low, ranging between 15 µM in the glacial and 21 µM in the non-glacial lake (Table 1). Enumeration of the microbial community revealed orders of magnitude higher abundances of non-phototrophic and phototrophic pro- and eukaryotes in the non-glacial as compared to the glacial lake (Table 1).

**Table 1.** Brief description of environmental parameters of the Lake Alexandra David Néel (ADN) with no contemporary influence by glacial melt and the glacial influenced Lake Ampère (AMP)

|  | Lake ADN | Lake AMP |
| --- | --- | --- |
|  | Non-glacial | Glacial |
| Dissolved iron, DFe (nM) | 113 | 964 |
| Phosphate, PO<sub>4</sub><sup>3-</sup> (µM) | 0.05 | 0.16 |
| Nitrate, NO<sub>3</sub><sup>-</sup> (µM) | 0.50 | 3.27 |
| Silicic acid, Si(OH)<sub>4</sub> (µM) | 41.9 | 31.9 |
| DOC (µM) * | 21 | 15 |
| Non phototrophic prokaryotes (cells mL<sup>-1</sup>) | 513747 | 7354 |
| Cyanobacteria (cells mL<sup>-1</sup>) | 62600 | 146 |
| Pico/nano-eukaryotes (cells mL<sup>-1</sup>) | 11300 | 59 |

\*data obtained from Thoppil et al. (2025), no DOC data available for the sampling carried out for the present study.

In line with flow cytometric observations, cyanobacteria, dominated by *Synechococcus* (closest relative WH5701) had also high contributions to total prokaryotic sequences (35% in the < 0.8 µm and 60% in the > 0.8 µm size fraction) in the non-glacial lake only (Supporting Information Fig. S2). The higher cell abundances of phototrophic and non-phototrophic microbes indicate biological activity in the non-glacial lake. The presence of glacial flour, only observed in the glacial lake, likely leads to a reduction in light penetration inhibiting autotrophic activity (Fig. 1). The lack of terrestrial organic matter input further constrains microbial heterotrophic metabolism. These different environmental settings could influence the quantity and composition of the colloids collected for the incubation experiments. While these colloids are predominantly clay minerals in both lakes, those originating from the non-glacial lake could be associated with an organic component of phytoplankton origin.

### Microbial growth in response to colloid additions

We calculated the final concentrations of non-glacial (46 nM, 147 nM and 462 nM) and glacial colloidal Fe concentrations (260 nM, 841 nM and 2637 nM) in our incubations based on the DFe concentrations in the respective RUF and the dilution factors. Our aim was to obtain the same final concentrations for both types of colloids and Fe-EDTA (10 nM, 32 nM and 100 nM). As it was not possible to measure the DFe concentrations in the RUF prior to the incubations, we determined the dilution factors based on the DFe concentrations in the RUF obtained from the same lakes during a previous study (Thoppil et al. 2025). The unexpected high final concentrations of colloidal Fe are most likely due to a higher efficiency of the ultrafiltration carried out in the present study. This experimental feature does not affect the interpretation of our results, as a main objective of the present study was to determine the microbial responses to dose additions of different types of colloids. The abundance of non-phototrophic prokaryotes in the < 1 µm filtered seawater collected at the HNLC site was 2.91 x 10^5^ cells mL^-1^, corresponding to 68% of unfiltered seawater (4.29 x 10^5^ cells mL^-1^). In the 3 unamended incubations, prokaryotic cell abundances increased by 1.46-fold during the 4 days (Fig. 2), reaching values similar to those observed in the initially unfiltered water (4.26 ± 0.12x 10^5^ cells mL^-1^; mean of triplicate incubations ± SD). The fold-changes of prokaryotic abundances in the incubations amended with Fe-EDTA were 1.45, 1.43, and 1.47 for 10 nM, 32 nM, 100 nM, respectively, and thus similar to those in the unamended incubations.

**Fig. 2.**
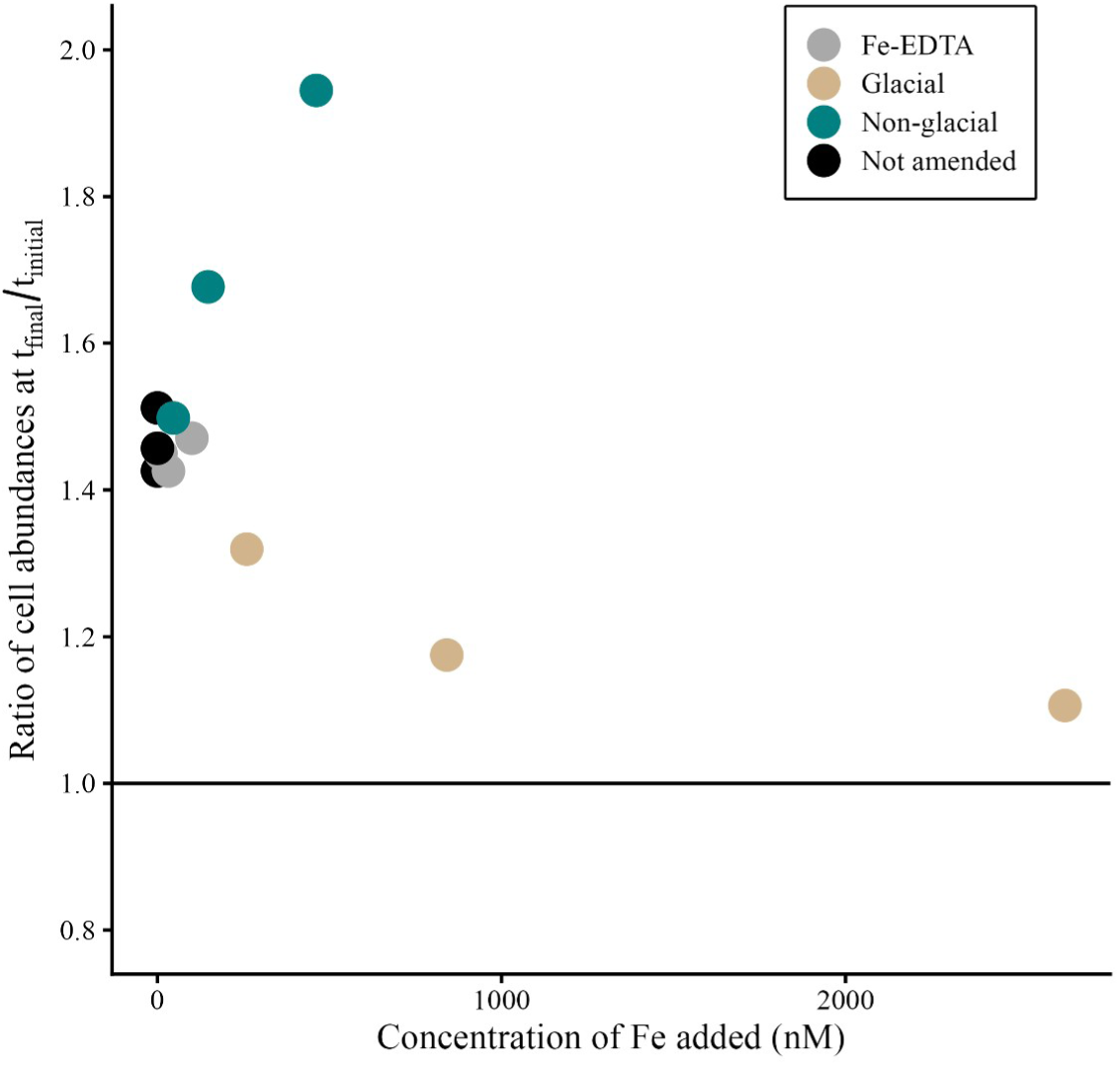
Relative increases in prokaryotic abundances after 4 days of incubation in the dark as compared to the initial time point. Fold-changes are shown as a function of Fe added in the form of non-glacial or glacial colloids, or as Fe-EDTA. Concentrations of Fe in the Fe-EDTA treatment were 0 nM, 10 nM, 32 nM and 100 nM. Black dots illustrate fold changes in unamended incubations.

Dose addition of colloids from the non-glacial lake resulted in an increasing trend in prokaryotic abundances, with fold-changes ranging from 1.50 to 1.94 (Fig. 2). By contrast, increasing additions of colloids from the glacial lake led to a decreasing trend in the fold-changes (range 1.32 to 1.11).

Heterotrophic nanoflagellates (HNF) increased with additions of non-glacial colloids by up to 8-fold, while abundances remained unchanged in the control and the glacial colloid amended incubations after 4 days (Supporting Information Fig. S3).

### Prokaryotic transcriptional response to dose addition of colloids

To further investigate the response of prokaryotes to the addition of Fe-rich colloids or Fe-EDTA in a dose-dependent manner we aimed to identify increasing trends in gene expression. A trend was defined as a 5% increase of a given metric (transcripts per million (TPM), transcripts per liter (TPL), and transcripts per active cell (TPC)) between increasing additions of colloids (see Material and Methods). The number of transcripts with an increasing trend was substantially higher in the incubations amended with glacial and non-glacial colloids as compared to those with Fe-EDTA (Fig. 3; Table S3; Table S4 to Table S10).

**Fig. 3.**
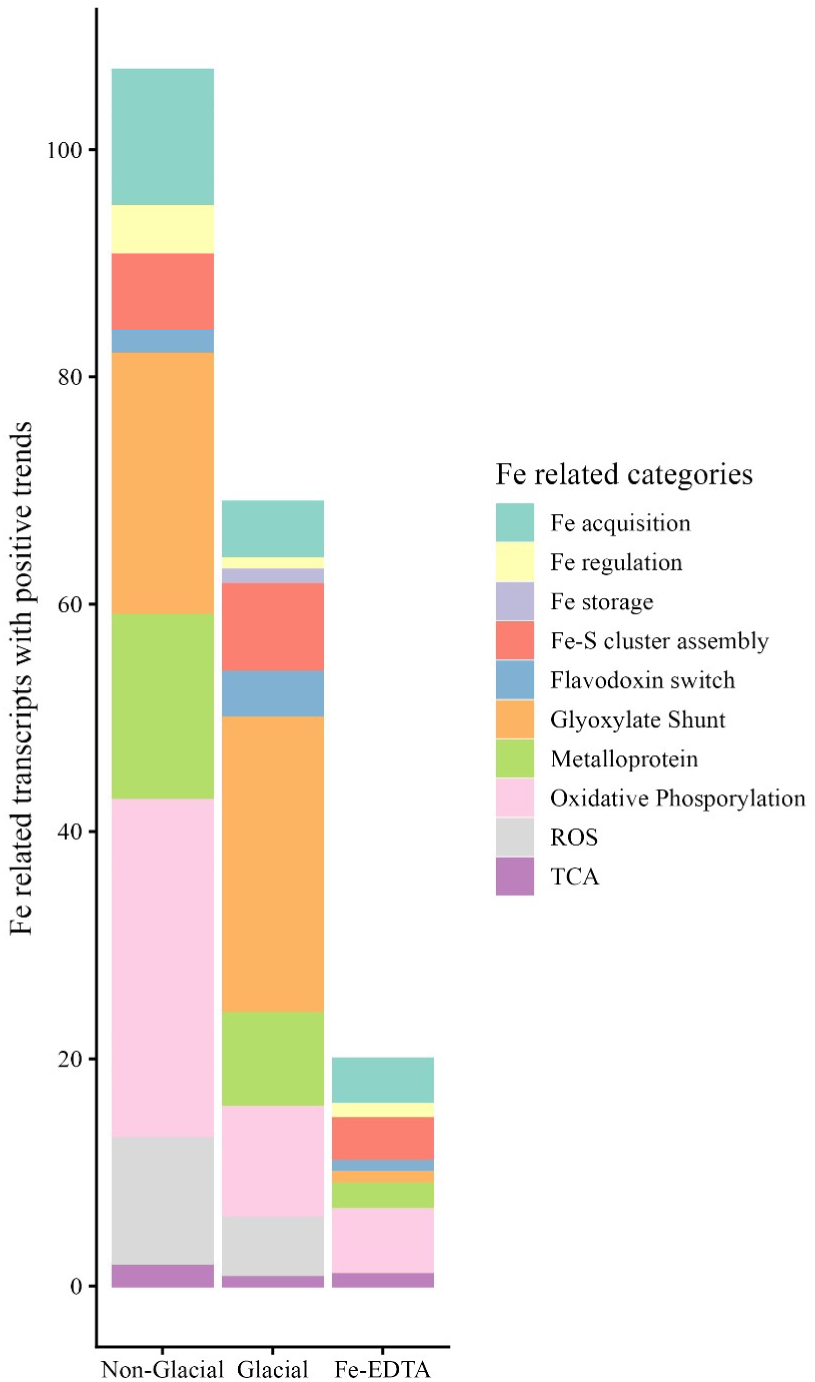
Category and number of Fe-related transcripts for which an increasing trend was observed in the incubations amended with glacial and non-glacial colloids and Fe-EDTA. Detailed information is provided in Table S13.

We then focused on Fe-related pathways, including Fe transport, regulation and storage, Fe metalloproteins and central carbon metabolism (detailed in Table S2), as Fe is a potentially limiting element provided by the colloids. We asked whether transcripts related to Fe had a statistically higher probability to show an increasing trend as compared to transcripts belonging to other categories. Using a Chi-square test, we observed significant results for non-glacial and glacial colloid amended incubations for all three metrics (Table 2, Table S12). By contrast, no significant results were obtained for the incubations amended with Fe-EDTA.

**Table 2.** Chi-square test results for increasing trends in Fe-related transcripts in incubations amended with non-glacial or glacial colloids, or Fe-EDTA. TPM-Transcripts per million. TPC – Transcripts per active cell. TPL-Transcripts per liter. Input data for the chi-square test are provided in Table S12. Trends within the TPL for the Fe-EDTA could not be calculated due to missing TPL values.

| Treatment | Metric | X-squared | P-value | Significance |
| --- | --- | --- | --- | --- |
| Non-glacial | TPM | 5.88 | 0.0152 | * |
|  | TPC | 10.15 | 0.0014 | ** |
|  | TPL | 5.36 | 0.0206 | * |
| Glacial | TPM | 4.75 | 0.0292 | * |
|  | TPC | 8.41 | 0.0037 | ** |
|  | TPL | 5.64 | 0.0175 | * |
| Fe-EDTA | TPM | 0.52 | 0.4687 | - |
|  | TPC | 0.36 | 0.5467 | - |

Fe-related transcripts involved in the glyoxylate shunt, oxidative phosphorylation, and metalloproteins contributed most to those with an increasing trend (Fig. 3). Transcripts of isocitrate lyase and malate synthase, key genes of the glyoxylate shunt, revealed a trend in incubations amended with either type of colloid (Fig. 4a). These transcripts represented an overall small portion of the total number of transcripts assigned to the respective genes, and they were taxonomically affiliated with *Burkholderiaceae*, *Flavobacteriaceae* and *Pseudomonadaceae* (Table S13) (Supporting Information Fig. S4). A trend was observed for only a few transcripts related to Fe and siderophore transport, such as *pir*A and *pvu*, and for several transcripts of the flavodoxin/ferrodoxin switch. Transcripts belonging to reactive oxygen species (ROS) were dominated by superoxide dismutase (Fe-Mn family) taxonomically affiliated to several families, including *Porticoccaceae*, *Flavobacteriaceae* and *Burkholderiaceae* (Table S13). Diverse transcripts for metalloproteins presented a trend, and among them the catechol 2,3- dioxygenase, taxonomically affiliated with *Burkholderiaceae,* was exclusively observed in the non-glacial colloid amended incubations (Table S13). Two pathways involved in xenobiotic degradation presented high transcription in the non-glacial incubation (Fig. 4b): benzene degradation (complete pathway) and the catechol meta-cleavage pathway (80% pathway completion) to which the catechol 2,3-dioxygenase belongs (Supporting Information Fig. S5).

**Fig. 4.**
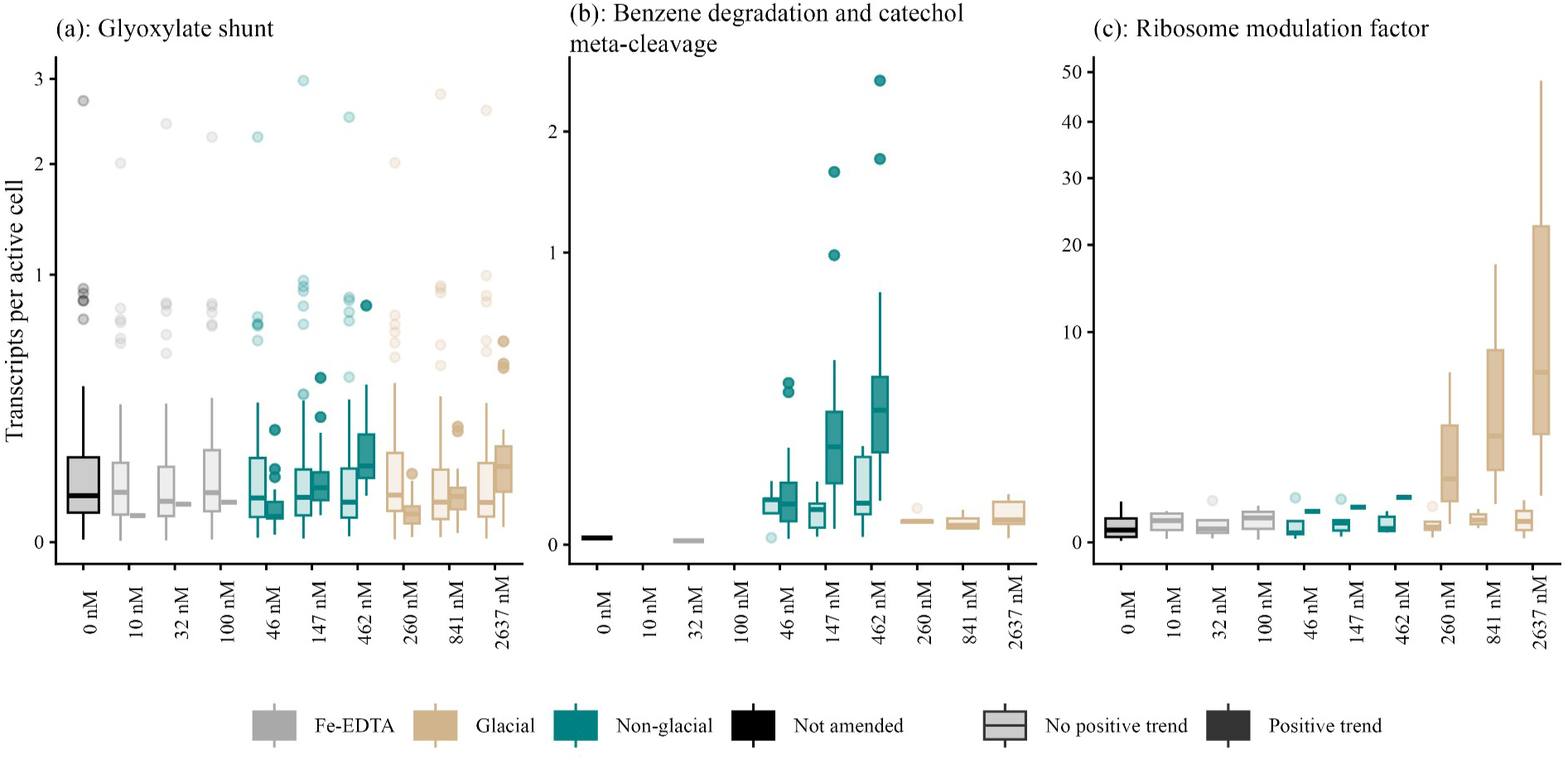
Transcripts per active cell in incubations amended with increasing quantities of Fe EDTA, non-glacial or glacial colloids. Transcripts are normalized to single copy genes. Median of transcripts with a 5% increase between sequential additions are shown in darker colors and transcripts with no increase are shown in lighter colors. (a) Pooled transcripts of the isocitrate lyase (K01637) and malate synthase (K01638) involved in the glyoxylate shunt. (b) Pooled transcripts of the pathways of benzene degradation and the catechol meta-cleavage pathway (detailed in Fig. S5). (c) Transcripts of the ribosome modulation factor (K03812).

Among the transcripts not directly related to Fe we observed an increasing trend and high expression levels for the ribosome modulation factor (Fig. 4c) and the hibernation promoting factor (Table S7 to Table S9) in the glacial colloid amended incubations. These indicators for stress induced ribosome inactivation had a lower transcription in the non-glacial incubations (Fig. 4c).

## Discussion

Our experiments revealed clear, does-dependent trends in both bulk and transcriptional responses of a prokaryotic community from the HNLC Southern Ocean to dose additions of Fe-rich colloid additions. These responses highlighted both shared and distinct effects of colloids from different origins on microbial metabolism and growth. The trace element Fe, a major constraint for microbial activity in the Southern Ocean, regulates heterotrophic metabolism in a complex interplay with the availability of organic carbon (C) (Fourquez et al. 2014; Koedooder et al. 2018). Within prokaryotic cells, the Fe and C cycles are tightly linked in the TCA cycle and electron transport system of the respiratory chain, a relationship that can lead to Fe- or Fe and C co-limitation (Church et al. 2000; Obernosterer et al. 2015; Baltar et al. 2018; Manck et al. 2020). Co-limitation arises when either both elements are insufficient to meet prokaryotic demands or when the scarcity of one restricts the utilization of the other (Saito et al. 2008). In our study, the latter scenario likely applied; prokaryotes could not utilize additional Fe sources when bioavailable C was limiting, and cellular Fe requirements had to be met to process available C. These insights help interpret prokaryotic responses to colloids of different origins in our incubations. Our results show that additions of Fe-EDTA, a highly labile Fe source, did not stimulate prokaryotic growth, suggesting that C, rather than Fe, was the limiting factor. In contrast, non-glacial colloids enhanced prokaryotic growth, indicating that they provided both Fe and C. The reduced growth of prokaryotes in the presence of glacial as compared to non-glacial colloids is, at first glance, puzzling. However, our dose-dependent transcriptional responses offer insights into the potential underlying mechanisms.

### The transcriptional profile reveals Fe-limitation

The transcriptional response of several Fe-related genes was independent of colloid origin, indicating Fe-limited prokaryotic communities in both incubations across all three metrics (TPM, TPL, TPC). Although transcripts for siderophore transporters (e.g. *pir*A), secondary metabolites produced under Fe-limitation (Andrews et al. 2003), were overall low in abundance, they exhibited a positive trend with increasing colloid addition. The flavodoxin/ferrodoxin switch is widely used as a marker of Fe-limitation in phytoplankton (Kolber et al. 1994), but the same substitution can also occur in metabolic pathways (e.g. respiratory chain) of heterotrophic microbes. However, the most pronounced transcriptional response was observed for genes involved in the glyoxylate shunt, a shortcut in the TCA cycle. This pathway allows cells to conserve C while aligning the production of reducing power with the demand of the Fe-limited electron transfer system chain (Fourquez et al. 2014; Koedooder et al. 2018). Previous studies have shown that the glyoxylate shunt is upregulated in Fe-limited prokaryotes as compared to non-limited sites (Debeljak et al. 2019; Sun et al. 2020; Manck et al. 2024) with expression levels scaling with the degree of Fe-limitation (Beier et al. 2015). Unexpectedly, transcripts for this pathway increased with colloid addition, despite both colloid types containing large amounts of Fe. These observations raise the question of the mechanisms drive this Fe-limitation-type response.

We propose two non-exclusive mechanisms that may explain the reduction of labile Fe in our incubations, regardless of colloid origin. These are the scavenging of labile Fe by colloidal particles and Fe uptake by prokaryotic activity occurring over the 4-day incubation period. From an operational perspective, the colloids used in this study were part of the dissolved phase (< 0.2 µm), which contains a diverse range of chemical Fe species. Geochemical analyses of the particulate phase (> 0.2 µm) from the two source lakes indicate that it is dominated by secondary aluminosilicate minerals (i.e. smectite) forming a continuum of particle sizes with similar chemical compositions (S. Blain, in preparation). Given the high clay content in both lakes, the colloids likely represent an extension of this continuum and thus contain similar material. When mixed with seawater such as in estuaries or, in our case, incubation bottles clay flocculation occurs, producing larger particles that may eventually sink. This process is likely dependent on particle concentration. Additionally, colloids have a high surface-to-volume ratio and a strong binding capacity for trace elements (Honeyman and Santschi 1989). Through this scavenging mechanism, labile Fe initially present in the seawater used for our incubations may have become biologically unavailable, leading to dose-dependent Fe depletion.

A second possible explanation is that prokaryotes rapidly utilize a fraction of the bioavailable Fe during the initial hours to days of the experiment. This aligns with observations from a previous study in which Fe-rich suspended marine particles were incubated with Southern Ocean microbial communities (Debeljak et al. 2021). In that experiment, cell-specific heterotrophic production in particle-amended treatments was approximately fourfold higher than in unamended controls at the start. However, this activity rapidly declined over the first 4 days, eventually matching control values (Debeljak et al. 2021). This transition phase was followed by a steady-state phase, during which prokaryotes processed the remaining particulate Fe (Debeljak et al. 2021). Thus, the rapid utilization of bioavailable Fe, potentially leached from colloidal matter, may have created a dose-dependent Fe-limited environment for the microbial community.

### Difference between glacial and non-glacial colloids

Despite shared transcriptional responses indicating Fe limitation, non-glacial and glacial colloids produced opposing patterns of prokaryotic growth, raising questions about the underlying mechanisms. In the case of glacial colloids, both bulk and transcriptional data suggest inhibition of prokaryotic activity. For example, the high abundance of transcripts of the ribosome modulation factor (RMF) indicates that cells were primed to inactivate their ribosomes and conserve energy (Yoshida et al. 2004; Njenga et al. 2023). However, this response contrasts with a previous study in which glacial colloids stimulated the growth of specific prokaryotic taxa, resulting in overall higher prokaryotic abundance as compared to the unamended control (Thoppil et al. 2025). In that study, incubations were conducted over a longer period (10 days) using a starting microbial community from a coastal site near the glacier meltwater outlet (Baie de la Table fjord, Kerguelen Island). This community may have been better adapted to utilize Fe from glacial colloids, as evidenced by the increased relative abundance of genes involved in the synthesis and transport of siderophores such as pyoverdine and vibrioferrin. Additionally, the higher concentrations of labile C in coastal seawater may also explain why prokaryotes were able to utilize Fe from glacial colloids (Thoppil et al. 2025).

In the case of non-glacial colloids, the dose-dependent increase in prokaryotic abundance suggests that these colloids provide both C and Fe. This idea could be further supported by the concurrent increase in heterotrophic nanoflagellates (HNF), which can utilize organic colloids as an alternative carbon source (Sherr 1988; Tranvik et al. 1993). If the increase in HNF abundance is due to the ingestion of prokaryotic cells, the stimulatory effect of non-glacial colloids on prokaryotic growth may be even more pronounced than observed. Additionally, we detected increased transcription of genes involved in the catechol meta-cleavage pathway, a major pathway for aromatic compound degradation, as well as benzene degradation genes (Supporting Information Fig. S5). The high expression levels of these genes suggest that aromatic compounds may serve as a carbon source provided by the colloids. In this pathway, catechol 2,3-dioxygenase cleaves the aromatic ring of catechol, producing 2-oxopent-4-enoate, which is subsequently metabolized to pyruvate and acetaldehyde, and ultimately to acetyl-CoA. Acetyl-CoA then enters the TCA cycle for energy production (Supporting Information Fig. S5). Notably, all these transcripts were exclusively affiliated with *Burkholderiaceae*, consistent with the known metabolic versatility of this bacterial family in degrading aromatic compounds (Pérez-Pantoja et al. 2012; Morya et al. 2020).

Benzene and other volatile organic compounds (VOCs), were long thought to originate solely from anthropogenic activities, but recent studies have identified biological sources in marine systems (Rocco et al. 2021). In freshwater environments, cyanobacteria have been shown to produce benzene, with nitrogen or phosphorus limitation as potential drivers (Fang et al. 2025a; b). Although we lack direct evidence, the relatively high cell counts of cyanobacteria and the high contribution of *Synechococcus* (clade WH5701) to total prokaryotic sequences in the non-glacial lake suggest this microbial group a potential candidate for VOC biosynthesis. This idea warrants further investigation. VOCs are insoluble and have been shown to bind with mineral clay particles (Chen et al. 2023), suggesting that benzene could be part of the colloidal fraction collected in the non-glacial lake. Regardless of the exact nature of the organic substrate provided by the colloids, prokaryotes had to metabolize C to sustain their enhanced growth. Our interpretation that Fe-limitation in this scenario is induced by unmet Fe-requirements is further supported by the observation that *Burkholderiaceae*, the taxa associated with transcripts of the catechol meta-cleavage pathway, were also responsible for most of the glyoxylate shunt transcripts, which exhibited a similar trend.

Our study demonstrates that dose-dependent additions of limiting nutrients in incubation experiments can elicit quantitative transcriptional responses, thereby providing mechanistic insights into observed bulk responses. By combining dose-addition experiments with metatranscriptomics we leveraged prokaryotes as ‘biosensors’ to uncover novel insights into the compounds and pathways involved in processing non-glacial origin colloids. The transcriptional activity point to differences in the chemical composition of the two colloid types, insights that chemical analyses alone cannot currently provide. Our results, obtained from incubations with prokaryotes from the HNLC Southern Ocean, highlight the critical role of bioavailable C in shaping microbial responses to Fe-rich colloids. Organic substrates alleviate C limitation and supply the energy required for siderophore production which is likely essential for accessing Fe bound in glacial colloids. In a parallel study, we observed a positive response of both prokaryotes and phytoplankton to glacial colloids when co-incubated (Thoppil et al. in prep).

This suggests that phytoplankton may facilitate prokaryotic access to Fe in colloids by mitigating C limitation. Such microbial reworking of glacial Fe could create positive feedbacks on phytoplankton growth, making microbial interactions key for unlocking glacial Fe as a novel source for microbial productivity in the Southern Ocean.

## Acknowledgements

We thank the captain and the crew of the R/V Marion Dufresne for their support during the MARGO cruise. We thank Patricia Medeiros (Univ. of Georgia, Athens USA) for providing the DOC concentration and Corentin Baudet (LEMAR, Brest, France) for providing the dissolved Fe concentration in surface waters at the HNLC site (MA02), Alizée Berthelot (LOMIC) for the amplicon sequencing analyses of the samples from the two lakes and Laurence Garzarek (SBR, Roscoff, France) for the precise identification of the *Synechococcus* ASV. The authors used Mistral AI to refine the writing style of this manuscript. All suggestions were reviewed and edited by the authors, who assume full responsibility for the final content. This work is a contribution to the projects MARGO (Matter of glacial origin and its fate in the ocean: a case study at Kerguelen; ANR-22-CE01-0004) and PIANO (Phytoplankton-prokaryote interactions in Antarctic environments influenced by ice sheet melting: the key role of iron; ANR-24-CE02-5808-01) and was further supported by the French Polar Institute (IPEV, project number 1184 and 1289), the French Oceanographic Fleet (FOF)(doi.org/10.17600/18002958), the CNRS LEFE-CYBER program, the CPER IDEAL and the University of Littoral Cote d’Opale (ULCO, poste ATER 1392). The outstanding assistance of the IPEV team throughout the project and in particular on Kerguelen Island made this work possible and is highly appreciated.

