## Supporting Information for "Dose-dependent growth and gene expression in marine prokaryotes in response to glacial and non-glacial iron-rich colloids from Kerguelen Island (Southern Ocean)"

2- CNRS, Sorbonne University, Laboratoire d’Océanographie Microbienne (LOMIC), 1 ave Pierre Fabre, 66650 Banyuls sur mer, France

Carla Pereira Garcia:

Urania Christaki :

Rhea Thoppil :

Audrey Guénueguès:

Philippe Catala:

Olivier Crispi :

Stephane Blain :

Ingrid Obernosterer:

*corresponding author

**This file includes:**

Supporting Information Material and Methods

Supporting Information Figures Fig S1, S2, S3, S4

**Other supplementary material for this manuscript includes the following:**

Supporting Information Tables 1-13 in excel format

**Inorganic nutrient analyses**

For nitrate (NO_3_^-^) and phosphate (PO_4_^3-^) lake water was collected by a 50 mL syringe that was directly connected to the spigot of the Niskin bottle. The samples were drawn through a 0.45 µm Uptidisc (Whatman) adapted for the syringe. The filtered samples were poisoned with mercuric chloride (HgCl_2_, 20 mg L^-1^ final concentration) and stored in the dark until analysed in the home lab. Concentrations of NO_3_^-^ and PO_4_^3-^ were determined with a segmented flow analyser (Skalar) equipped with a colorimetric detection using methods described previously (AMINOT A. and KEROUEL R. 2007)

**Dissolved organic carbon**

As described in Letourneau et al. (2021) concentrations of DOC from water samples were measured with a Shimadzu TOC-L_CPH_ analyzer using potassium hydrogen phthalate as a standard for the DOC calibration curve. Prior to and alongside sample analysis, both internal blanks and Milli-Q water blanks were run on the instrument. Analytical accuracy and precision were tested against the Consensus Reference Material (Hansell 2005) and were better than 5%.

**Dissolved Fe determination**

Lake samples (30 ml) were collected in trace metal clean low HDPE bottles and acidified with 30µL of HNO_3_ (Merck Suprapur grade). The samples were stored at room temperature until analysis in the laboratory a few months later. The dissolved Fe concentrations were determined by Q-ICPMS (Agilent). The accuracy (5%) was determined using the Standard Reference Material 1643 (NIST) containing trace elements in fresh water. Sample collection and analysis of dissolved iron at the HNLC site (MA02) was carried out as described in (Baudet et al. 2024).

**Enumeration of microbial community members**

For the enumeration of phototrophic and non-phototrophic cells in the two lakes, unfiltered lake water (4.32 mL and 1.44 mL, respectively) was ﬁxed with glutaraldehyde grade I 25% (1% ﬁnal concentration), incubated for 30 min at 4°C, and then stored at -20°C (roughly 1 week) and at -80°C until analysis in the home lab. For the enumeration of non-phototrophic cells and heterotrophic nanoflagellates (HNF) at the end of the incubation experiment subsamples (1.44 mL and 4.32 mL, respectively) were fixed with glutaraldehyde, incubated for 30 min at 4°C, shock frozen in liquid nitrogen and then stored at -80°C until analysis. Prior to flow cytometric analyses, samples were thawed at room temperature. Counts were performed on a FACSCanto II ﬂow cytometer (Becton Dickinson) equipped with 3 air-cooled lasers: blue (argon 488 nm), red (633 nm) and violet (407 nm). For the enumeration of non- autoﬂuorescent cells, mainly heterotrophic prokaryotes, cells were stained with SYBR Green I (Invitrogen – Molecular Probes) at 0.025% (vol/vol) ﬁnal concentration for 15 min at room temperature in the dark. Stained prokaryotic cells were discriminated and enumerated ac-cording to their right-angle light scatter (SSC) and green ﬂuorescence at 530/30 nm. Fluorescent beads (1.002 μm; Polysciences Europe) were systematically added to each analysed sample as an internal standard. The cell abundance was determined from the ﬂow rate, which was calculated with TruCount beads (BD biosciences). The enumeration of different autotrophic populations was performed in a second analysis, it was based on their scattering and fluorescence signals according to Marie et al. (2000). *Synechococcus* spp. was discriminated by its strong orange fluorescence (585 ± 21 nm), and pico- and nano-eukaryotes were discriminated by their scatter signals of red fluorescence (> 670 nm). Heterotrophic nanoglagellates were enumerated after staining with SYBR Green I (Invitrogen – Molecular Probes) at 0.025% (vol/vol) as described previously (Christaki et al. 2011).

**DNA extraction**

DNA extraction for all samples was performed using the DNeasy PowerWater Kit (Qiagen, Hilden, Germany), following the manufacturer’s protocol with minor modifications to optimize the lysis of the cells. In the first step in the manufacturer’s protocol, additionally to the PW1 solution, we added 66µl of lysozyme at 20mg/ml and incubated at 37˚C for 45 minutes, followed by the addition of 14 µl of proteinase K at 20mg/ml and incubation at 55˚C for 60 minutes. Final DNA concentrations were measured in the home laboratory using the QuantiFluor® DNA System.

**16S amplicon sequencing**

For the PCR amplification of the two lake samples we use the following conditions: an initial denaturation step of 95 ◦C for 3 min, followed by 30 cycles of denaturation at 95°C for 45 s, annealing at 50°C for 45 s, and extension at 68°C for 90 s, and a final elongation step at 68°C for 5 min. Amplicon sequencing variants (ASVs) were obtained with DADA2 v1.38.0 (Callahan et al. 2016) based on the following parameters: truncLen=c(200,200), maxN=0, maxEE=c(2,2), truncQ=2, rm.phix=TRUE, trimLeft = c(10,10). The pipeline consisted of certain steps:  quality filtering and trimming, error-rate learning, dereplication and sample inference, paired-end read merging, sequence table construction, chimera removal and singletons removal. A mock community with known reference sequences of bacterial strains were evaluated to check the accuracy of DADA2 and the residual error rate of the analysis was 0%, ensuring that the ASVs identified by DADA2 present in the mock community exactly matched the reference sequences of expected bacterial strains.

**Normalization approaches:** To investigate the transcriptional responses in a detailed manner, we used different normalization approaches. Transcripts per million (TPM) is a metric commonly used to assess the relative abundance of the transcripts (Equation 1); it takes into account the gene length and the sequencing depth of the sample. The second metric, transcripts per liter (TPL), used in this study is to evaluate the absolute response of the community (Satinsky et al. 2013). This approach uses an internal standard added during the first step of the extraction to calculate the total transcripts of genes of interest in the sample (Equation 2). A more recent approach, proposed by (Salazar et al. 2019), uses ten single copy genes to normalize the number of transcripts by the transcriptionally active cells (Equation 3).

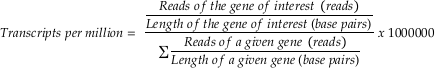

Equation 1: Calculation of TPM

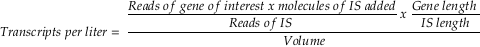

Equation 2: Calculation of TPL. IS: internal standard.

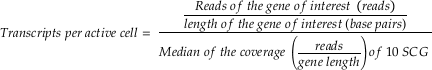

Equation 2: Calculation of the transcripts per active cell (TPAC). SCG: single copy genes.

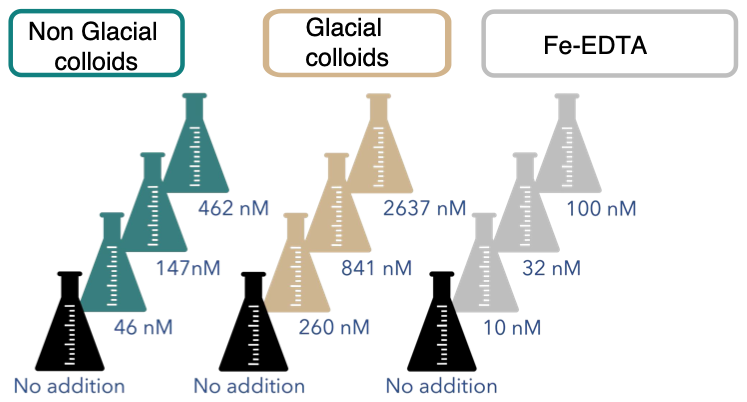

**Fig. S1.** Schematic illustration of the experimental set-up used in the present study. Non-glacial and glacial colloids, and Fe-EDTA were added to 1 µm filtered surface seawater from an HNLC site in the Southern Ocean. Final concentrations of Fe in the form of colloids and Fe-EDTA are given.

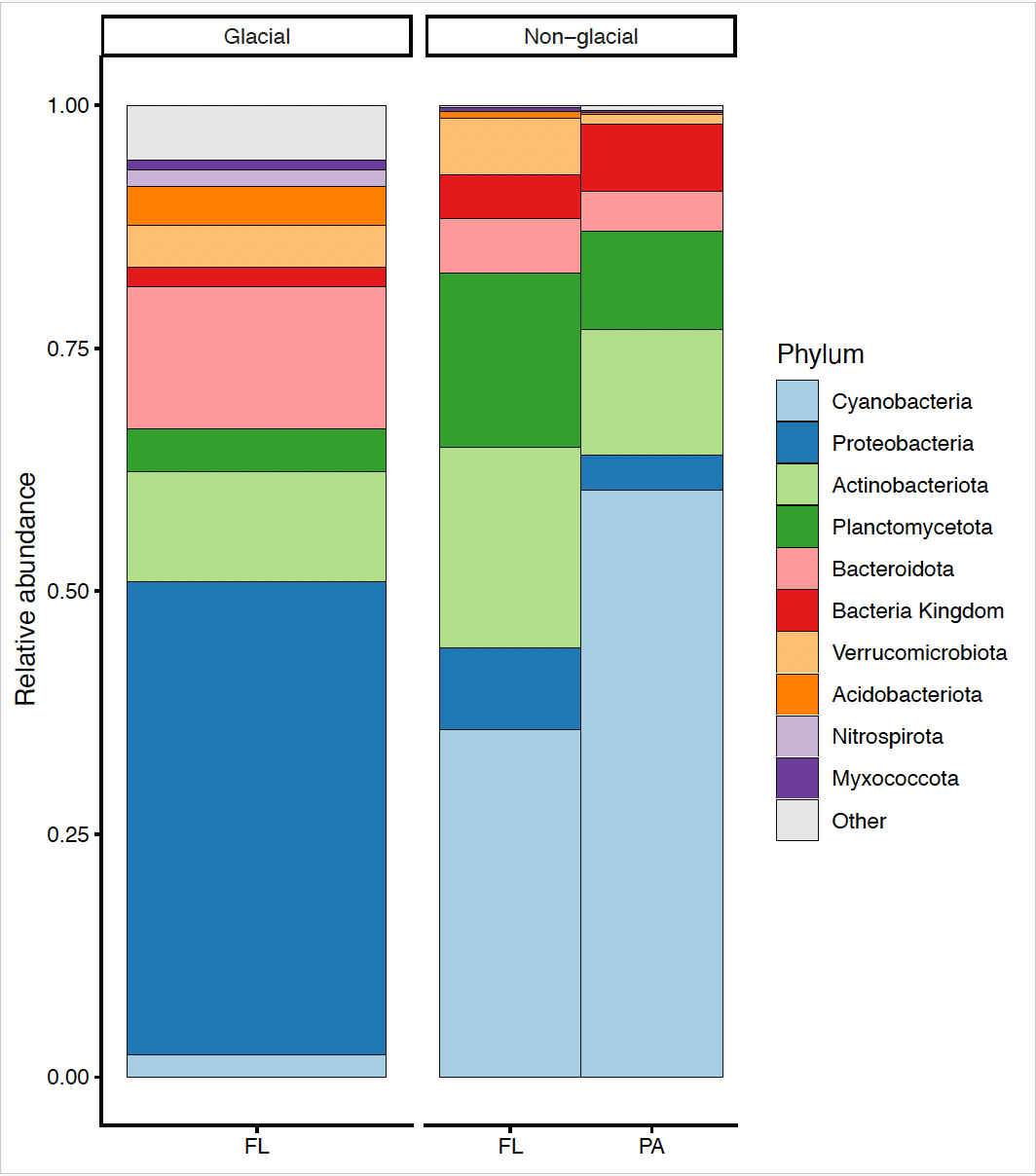

**Fig. S2**. Composition of prokaryotic communities in samples collected at the surface of lake Ampere (glacial lake) and Lake Alexandra David Néel (non-glacial lake) based on 16S rRNA sequencing. Relative abundances are shown for the free-living (FL, < 0.8 µm size fraction) and the particle-attached fraction (PA, > 0.8 µm size fraction). The particle-attached fraction of the glacial lake had to be excluded due to low DNA recovery after extraction.

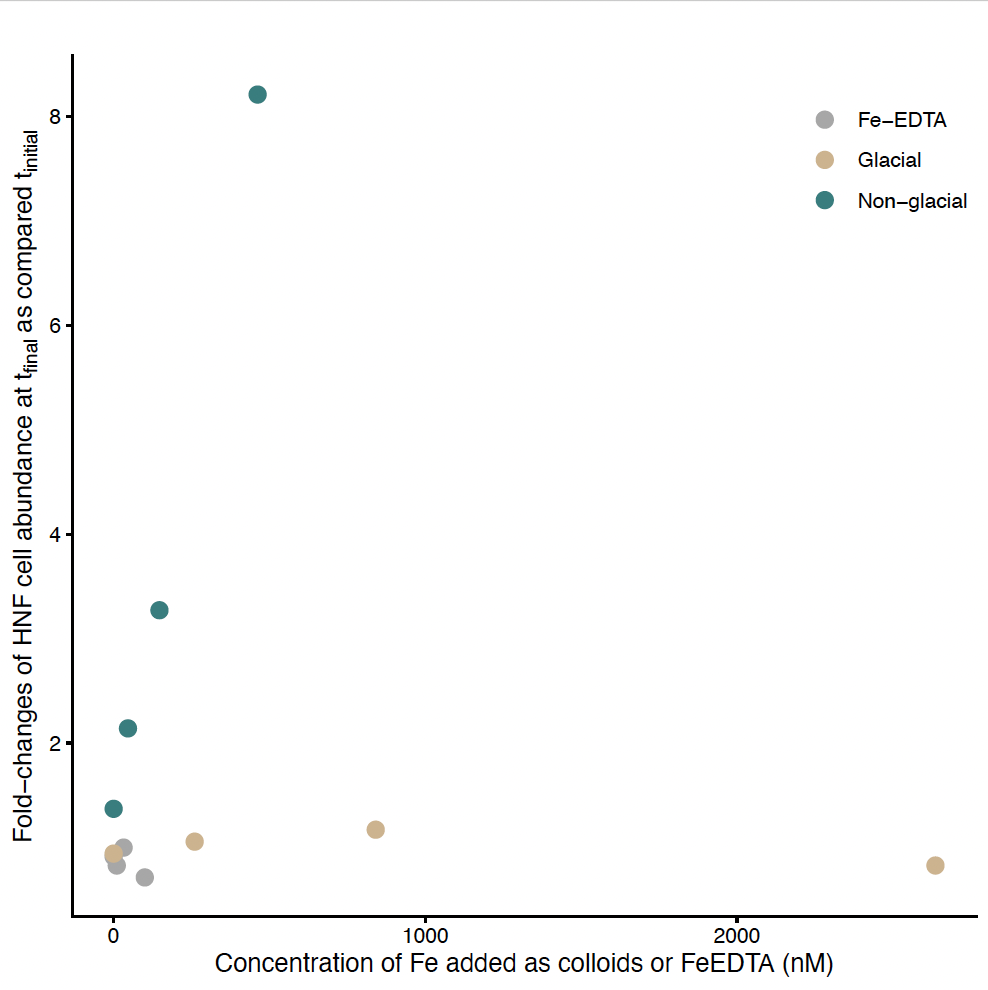

**Fig. S3.** Relative increases in heterotrophic nanoflagellate (HNF) abundances after 4 days of incubation as compared to the initial time point. Fold-changes are shown as a function of Fe added in the form of non-glacial or glacial colloids, or as Fe-EDTA. Concentrations of Fe in the Fe-EDTA treatment were 0 nM, 10 nM, 32 nM and 100 nM.

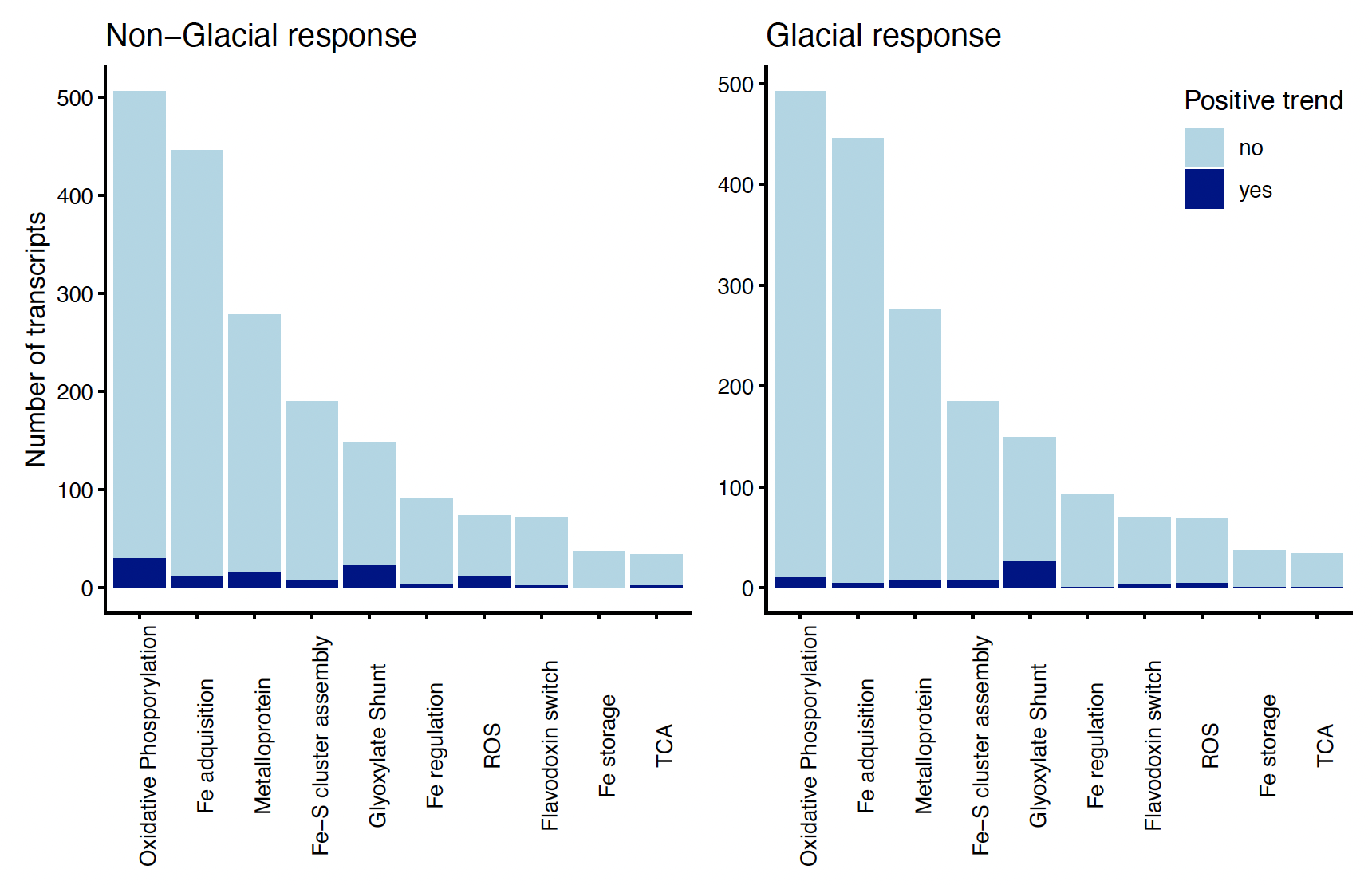

**Fig. S4**. Total number of Fe-related transcripts of a given category. Transcripts which show an increasing trend (yes) are represented in dark blue and transcripts without a trend (no) are represented in light blue.

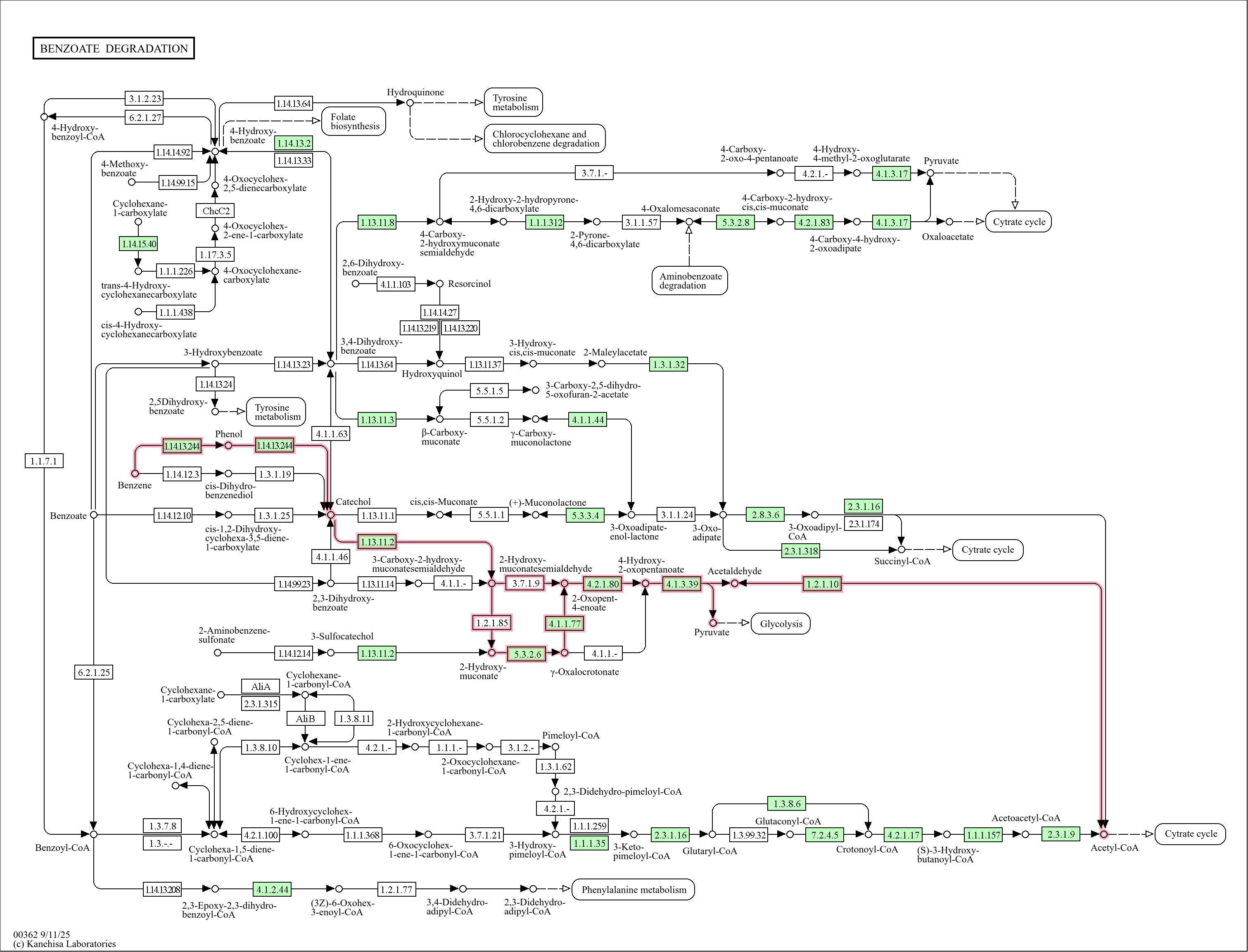

**Fig. S5**. Full Benzoate degradation pathway map from KEGG. Green boxes represent all the genes being transcribed in the non-glacial samples and the benzene degradation and catechol meta-cleavage pathways are highlighted in pink.
